# Genomic prediction for malting quality traits in practical barley breeding programs

**DOI:** 10.1101/2020.07.30.228007

**Authors:** Pernille Sarup, Vahid Edriss, Nanna Hellum Kristensen, Jens Due Jensen, Jihad Orabi, Ahmed Jahoor, Just Jensen

## Abstract

Genomic prediction can be advantageous in barley breeding for traits such as yield and malting quality to increase selection accuracy and minimize expensive phenotyping. In this paper, we investigate the possibilities of genomic selection for malting quality traits using a limited training population. The size of the training population is an important factor in determining the prediction accuracy of a trait. We investigated the potential for genomic prediction of malting quality within breeding cycles with leave one out (LOO) cross-validation, and across breeding cycles with leave set out (LSO) cross-validation. In addition, we investigated the effect of training population size on prediction accuracy by random two, four, and ten-fold cross-validation. The material used in this study was a population of 1329 spring barley lines from four breeding cycles. We found medium to high narrow sense heritabilities of the malting traits (0.31 to 0.65). Accuracies of predicting breeding values from LOO tests ranged from 0.6 to 0.9 making it worth the effort to use genomic prediction within breeding cycles. Accuracies from LSO tests ranged from 0.39 to 0.70 showing that genomic prediction across the breeding cycles were possible as well. Accuracy of prediction increased when the size of the training population increased. Therefore, prediction accuracy might be increased both within and across breeding cycle by increasing size of the training population

## Introduction

Barley (*Hordeum vulgare* L.) was one of first the cereals to be cultivated. European barley production comprised around 7 million hectares in 2019 (CoCeral Grain crop forecast September 2019). Up to 50% of this barley production is used as malting barley; the rest is primarily used for animal feed and human consumption (FAOSTAT). Measuring malting quality is an expensive and labor-intensive process and the high costs of obtaining these phenotypes limits their use in development of new barley cultivars with better malting quality (Gao et al. 2004).

Malting properties is an important quality trait of barley. It is essential for the brewing and distilling industry, and therefore barley cultivars with high malting quality gain higher prices than cultivars with low malt quality. To start the malting process, the grains are steeped in water, which induces a controlled germination. Hydrolytic enzymes break down the endosperm cell walls making the starch, proteins and lipids stored in the cells available for hydrolyses by malting enzymes (Bamforth 2009). These nutrients are used as building material and energy source for the seedling in a natural germination process. In brewing, the malt is milled and extracted in water; the resulting malt extract (wort) provides the nutrients and energy for the brewing yeast that is required for fermentation and production of alcohol. To maximize the sugar concentration available for the yeast, good malting barley should have high conversion rates from starch to fermentable sugars and high extract yield increases the amount of substrate available for fermentation. The protein content should be between 9,5 and 12,5% of dry weight and (1-3;1-4)-beta-glucan (BG) content should be low (Bamforth 2009). At least four enzymes (α-amylase (Alpha), β-amylase (Beta), limit dextrinase, and β-glucosidase) are involved in hydrolysis of starch (Delcour and Verschaeve 1987). Moreover, the prevalence of different isoforms of these enzymes during malting affects wort fermentability (Evans et al. 2005). The essential amino acids for yeast growth comes from degradation of grain proteins during fermentation; too high grain protein content on the other hand leads to a low malt extract and is, thus, not desirable (Qi et al. 2005). Filtration problems can be caused by complexes generated by disulphide bridges between hordeins and gel-forming proteins (Celus et al. 2006). Filtration problems can also be caused by high BG content leading to a highly viscous wort. High BG content can be a sign of insufficient degradation of cell walls. The remaining fragments of cell walls may hamper diffusion of enzymes in the germinating grain and therefore decrease yield of malt extract (Bamforth 2003). Thus, malting quality is composed of several interacting traits each controlled by many genetic and environmental factors and is therefore classical polygenetic quantitative traits (Hayes et al. 1993; Li et al. 2009).

Genomic prediction (GP) (Meuwissen et al. 2001; Goddard and Hayes 2007; Fernando et al. 2008) is a method that uses genotype information to predict the additive genetic value of individuals for specific traits. The genotype information comes from dense single nucleotide polymorphism (SNPs) spread across the whole genome. This method exploits linkage disequilibrium (LD) between QTLs and SNPs (Meuwissen et al. 2001) and assumes that the effect of chromosome segments will be consistent across genotypes. This requires sufficient marker density to cover the whole genome in order to ensure that all QTLs are in LD with at least one marker or marker haplotype. The use of high throughput genotyping technologies combined with decreasing genotyping costs provides dense markers set required for saturation of the genome, thus providing the possibility to widely apply GP in breeding programs. To implement genomic prediction, the first step is to split the population into two groups of a reference and a validation population. The reference population is the individuals with genotypic and phenotypic information and these individuals are used to estimate model parameters. For the validation population, phenotypes are withheld from the model and only their genotypic information is used in the prediction in order to emulate the situation, where lines in the breeding population only have genotypes but no phenotypes. The genetic value of the candidates is predicted using the estimates of model parameters from reference population. Then the accuracies of prediction in the validation population can be estimated by comparing predicted genetic value and actual phenotypic performance.

Classic barley breeding programs are mainly based on mean phenotypic performance of each potential line. In genomic selection-based breeding programs, we can select superior lines before we have their phenotypes. This means that the number of candidate lines to select from is no longer limited by the capacity for field trials or laboratory tests and can be increased resulting in much higher selection intensities. If prediction accuracies are high enough, the added selection intensity results in higher genetic gain per generation than using phenotypic selection and justify the added cost by genomic prediction compared with the traditional breeding programs. Previous simulation and empirical genomic prediction studies have shown that multiple factors affect prediction accuracy. These factors include the genetic architecture of the trait, marker density, genetic relationship between training and validation sets, composition of the training population in terms of relatedness of individuals, heritability of the trait, population size and LD (Hayes et al. 2009; Lorenz et al. 2011; Riedelsheimer and Melchinger 2013; Nielsen et al. 2016). Higher prediction accuracies have been reported in genomic prediction studies when the training population is increased and the marker densities are higher, and when there are close genetic relationships between the training and the validation populations (Lorenz et al. 2011). Therefore, the training population needs to be constantly updated to ensure a close genomic relationship between training and validation population.

In recent years, genomic prediction have become popular in cereal breeding (Heffner et al. 2009; Jannink et al. 2010; Nielsen et al. 2016; Cericola et al. 2017; Kristensen et al. 2019). A major advantage of genomic prediction is that candidate individuals are only genotyped and need not to be phenotyped at the selection stage. As selection of the seedling happens at an early stage, genomic prediction is cost effective and selection cycle time can be reduced which increase the genetic gain per time unit (Heffner et al. 2009).

The aim of this study was to investigate the accuracy of genomic prediction for malting quality traits in barley. This has already been assessed in Bhatta et al. (2020) for a small closed population (145 lines originating from crosses using 5 different parents). Whether the conclusions drawn in the study of Bhatta *et al.* is relevant for a large practical barley breeding population still remains an open question. We used a large spring barley population consisting of four breeding cycles from a commercial breeding program (1329 lines in total). To evaluate the potential for genomic selection in a practical breeding program, the effect of timing of the evaluation on prediction accuracy was investigated using three types of cross-validation.

1. Leave set out cross-validation (LSO) was used to estimate prediction accuracies, when evaluating new lines before phenotypes from any of the lines in a breeding cycle was obtained (predicting across breeding cycles).
2. Leave one out cross-validation (LOO) was used to estimate prediction accuracies when evaluating new lines after phenotypes from a subset of the lines in a breeding cycle was obtained (predicting within breeding cycles).
3. Random selected k-fold validation sets were used to investigate the effect of training population sizes on prediction accuracies.

## Materials and methods

### Plant material

All plant and genetic material were supplied by Nordic Seed A/S, a Danish breeding company, who also conducted all field trials and all malting quality analysis. The germplasm consisted of 1329 advanced spring malting barley lines from Nordic Seed. The lines were harvested in four different years (2013-2016). All lines were grown in three locations in Denmark: Skive (56°37’38.0”N 9°02’35.7”E), Dyngby (55°56’57.2”N 10°15’13.8”E) and Holeby (54°42’03.1”N 11°27’07.6”E). In each year-location lines were sown in three replicates. The fields were divided in trials consisting of smaller plots, and each trial was a randomized block. Two control lines were sown in three replications in each trial, 412 lines were sown in all locations in two consecutive years 917 lines were sown in all locations in one year only. The size of each plot was 5.5 × 1.5 m (8.25 m^2^) and sample seeds harvested from each plot were used for measuring malting quality.

### Phenotyping

Seed samples were first sorted individually by a SORTIMAT (Baumann Saatzuchtbedarf) instrument. Seeds larger than 2.5 mm were used for micro malting in 40 g per sample. Batches of 250 samples were malted using an automatic micro-malting system. The steps in the malting process were adjusted to the harvest year in the beginning of the season. The basic steps of the process were 4 h of steeping, 18 h of air flow (for resting), 4 h of steeping after which seeds germinated for 5.5 days at 16 °C and relative humidity at 100%. Samples were kilned in a heating chamber for 14 h at 45 °C, followed by 65 °C for 4 h and ending with 6h at 80 °C. The malted barley samples were milled for further analysis.

After milling, 5.5 g of grounded flour was poured in a plastic container for malting experiment. From the same flour, two extra samples were taken for measuring the moisture content. A total of 44 ml of distilled water was added to the malt sample container. The samples were then mashed in batches of 47 samples. The malt sample containers were placed in a large water bath for 2 hours to dissolve the milled powder in water. During this time, the samples were shaken. After mashing, the samples were poured in a filter, and the wort was used to measure malting quality traits. Filtering speed was scored by measuring the height of the liquid surface in the glass 20 minutes after filtering was begun. Clearness of the wort was evaluated visually at this step by scoring each filtrate from 1-3, where 1 is clear and 3 is opaque. After filtration, the wort samples were separated in two parts and all wort phenotypes were obtained according to the Analytica–EBC 2004 manual. One sample of 25 ml of wort used for viscosity (Analytical–EBC 8.4) and extract yield (Analytical–EBC 8.3). Second sample of 3-4 ml of wort was used for beta-glucan (Analytical–EBC 8.13.1), free amino nitrogen (FAN) (Analytical–EBC 8.10) and color measurements (Analytical–EBC 8.5).

### Genotyping

DNA extraction was carried out from individuals of each the 1329 advanced lines. DNA from three seedlings (two-weeks old) were bulked for each line. A modified cetyl trimetheyl ammonium bromide (CTAB) procedure was used to extract DNA (Saghai-Maroof et al. 1984). All lines were genotyped by the Illumina iSelect9K barley chip containing a total of 7865 markers. The genotyping was performed by TraitGenetics (Gatersleben, Germany). After filtering a total of 4,710 polymorphic markers were obtained from the 9K barley chip. Marker editing was done based on minor allele frequency more than 1% and missing markers less than 20%.

### Statistical analysis

Principal component analysis (PCA) and a heatmap were produced from the relationship matrix to check, if any clear sub-population exists in the population. The R software was used for both the heatmap and the PCA (prcomp command).

### GBLUP

The genomic relationship matrix was calculated using principles described by VanRaden (VanRaden 2008) and was used in the genomic prediction model fitted using DMU (Madsen & Jensen, 2013). Linear mixed models with fixed and random effects were used to predict genomic breeding value:

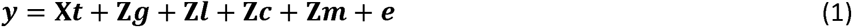

Where *y* was the vector of phenotype observations, ***t*** was the vector of fixed effects. The fixed effect was year×location×trial. **Z** was the design matrix for random factors, ***g*** was the vector of additive genomic breeding values 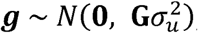, **G** was the genomic relationship matrix, ***l*** was the vector of line effects not explained by markers 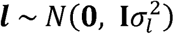, **I** was the identity matrix, ***c*** was the vector of year×location×line effects 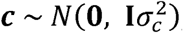, ***m*** was the vector for malting and mash batch effects 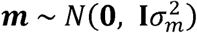, and ***e*** was the vector of random residuals effect 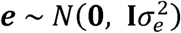. The estimates of narrow sense heritability (h^2^) of line means was calculated as 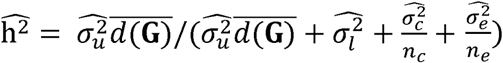 where *n*_*c*_ and *n*_*e*_ was the average number of observations per line for G×E and residuals respectively, 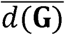 was the average of diagonal elements in **G**.

In addition to the 7 single trait models (one for each trait) a 7-trait model including all traits was fitted to estimate genetic and phenotypic correlations among traits.

### Cross-validation

Three different types of cross-validation schemes were used to evaluate different aspects of predictive ability of genomic models for malting quality. These types were:

Leave one out (LOO) was used to evaluate the ability to predict the genomic value of a line from a breeding cycle, where other lines have own phenotypes. In this method, phenotypes of a single line were set to missing and its genomic value was predicted using the rest of the lines. Naturally, the relationship between the training population and the validation line are strongest and the training population is largest in the LOO cross-validation (CV) scheme. Thus, we expect the highest predictive ability here.

In breeding programs, every year, a new set of lines are developed and preferably should be predicted before phenotypes from the new breeding cycle can be obtained. To check, if the model was able to predict one cycle from rest of the cycles, the LSO (Leave Set Out) scheme was applied. In this scheme, every phenotypic observation coming from a single breeding cycle was set to missing and the genomic values for all the lines included in that breeding cycle were predicted. This CV-strategy tests, whether future sets of crosses can be predicted based on current and previous sets of crosses.

The last method was K-fold cross-validation. The lines were randomly divided into k-folds of equal size and one fold acted as validation set and the remaining lines as training set. The validation and training sets were the same for all traits. Two-, four- and tenfold were applied in this study each with ten replications. For example, in fourfold random cross-validation, 25% of the lines were assigned to the validation population and the remaining 75 % as training population. Thus, the fourfold cross-validation had the same size training population as the LSO cross-validation. The aim of this cross-validation using different folds was to evaluate the effect of training population size. If, for example, the fourfold cross-validation set of a specific trait had lower predictive ability than the tenfold cross-validation set of the same trait, this would indicate that the LSO predicting accuracy under estimate the prediction accuracy of a new breeding cycle based on a training set including all the four breeding cycles with phenotypic data.

Phenotypes were corrected for effects of location, year and trial for each line. Predictive ability (PA) was then calculated as the correlation between average of corrected phenotypic values for each line and the genomic predicted breeding value. The breeding value only represent the additive genetic variation in the phenotype and not *e.g*. the environmental variation or the genotype x environment interaction. Thus, the maximum correlation to the phenotypic values is the square root of the narrow sense heritability of line means. Thus, in order to evaluate, how well the models predict breeding values, we calculated the prediction accuracy by dividing predictive ability by the theoretical maximum, *i.e*. the square root of h^2^ of line means.

Venn diagrams was used to show the overlapping lines form the 200 lines that had the best genomic breeding values for a given trait in 2 groups of traits. Venn diagrams were plotted using online tools Venny 2.1 (http://bioinfogp.cnb.csic.es/tools/venny/).

## Results

### Phenotyping

Malting quality traits were measured on grains from several plots of 1329 advanced spring barely lines from four different breeding cycles (see Table 1). The phenotypes measured were extract yield, filtering speed, wort color, beta-glucan, viscosity, wort clearness and free amino nitrogen (Fig.1, Table 1). Residuals from the genomic models were normally distributed (figures in supplemental material S1). The phenotypic variance was lowest for extract yield and highest for beta-glucan with coefficient of variance equal to 1.6% and 62%, respectively.

**Table 1.** descriptive statistics of the malt quality traits.

| 236 Trait | Number of obs. | Average | SD | Min value | Max value |
| --- | --- | --- | --- | --- | --- |
| 237 Extract yield | 7525 | 82.2 | 1.77 | 63.43 | 92.39 |
| 238 Filtering speed | 8859 | 4.45 | 0.69 | 0.90 | 6.30 |
| 239 Wort color | 8942 | 5.73 | 0.96 | 3.30 | 10.58 |
| 240 Beta-glucan | 8808 | 200 | 144 | 50.0 | 1318 |
| 241 Viscosity | 7008 | 1.47 | 0.10 | 1.01 | 2.74 |
| 242 Wort clearness | 8948 | 1.15 | 0.42 | 1.00 | 3.00 |
| 243 Free amino nitrogen | 5670 | 187 | 49.2 | 51.2 | 336 |

**Fig.1.**
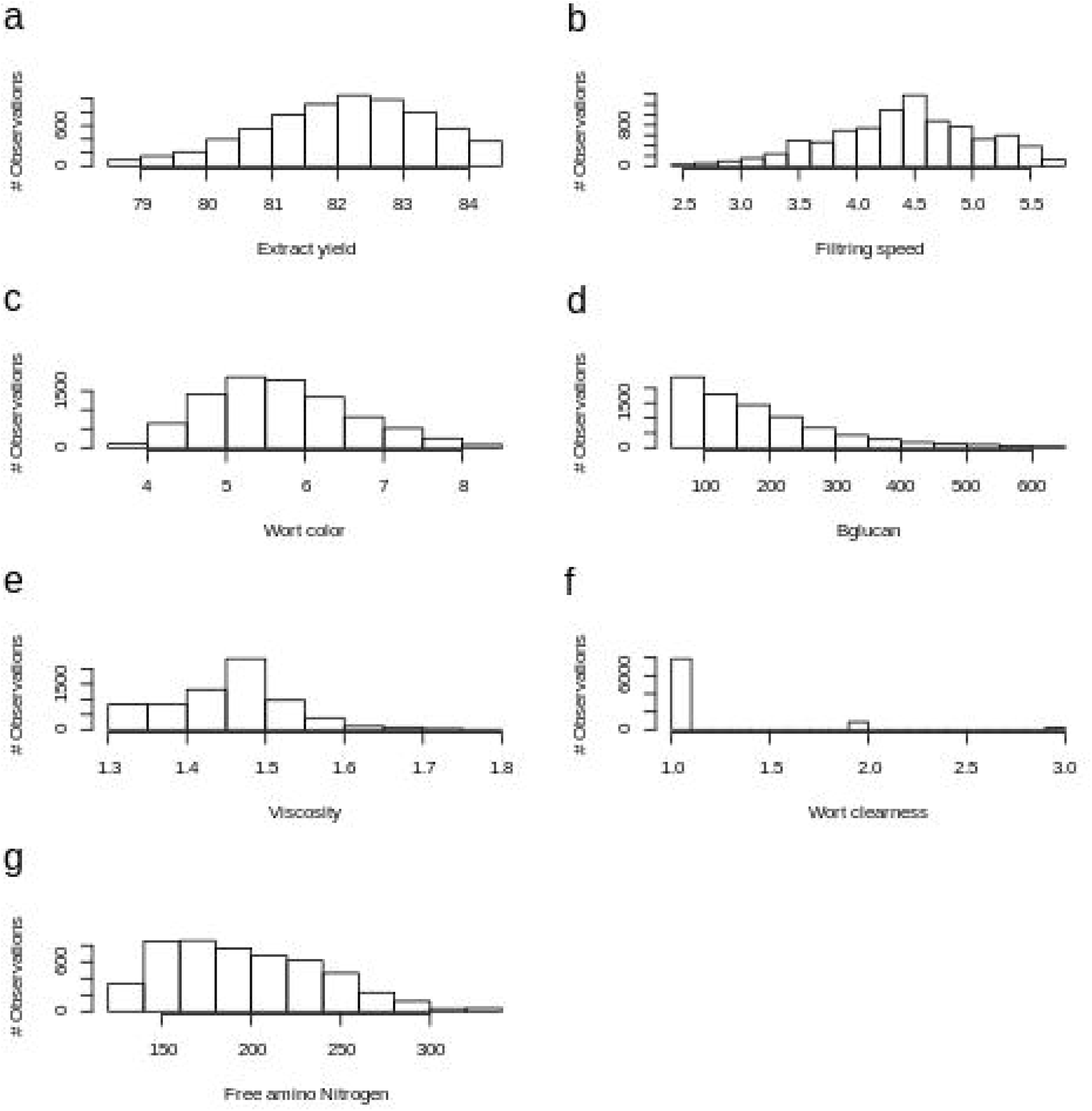
Phenotype distributions of malting quality traits.

### Genotyping

Genotyping of the 1329 barley lines were done with 9K Illumina Infinium BeadChip. A total number of 7,000 SNP markers were called. After filtering, the SNP markers based on minor allele frequency (MAF > 0.01) and missingness (< 0.2) 4,710 SNP markers were retained for analysis. Overall, the MAF and missing distribution of markers indicated high quality of the SNP marker dataset (Fig.2).

**Fig.2.**
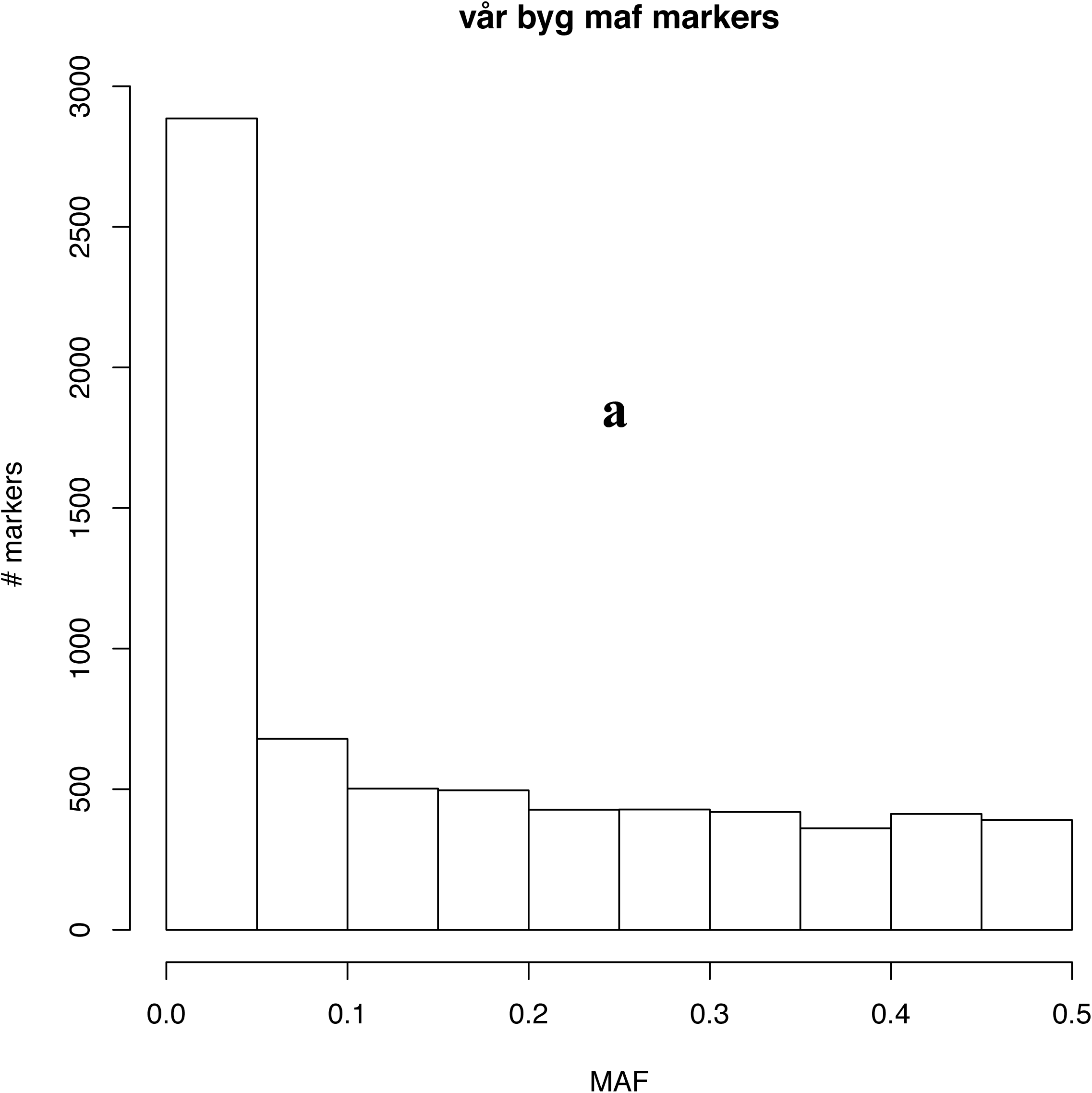
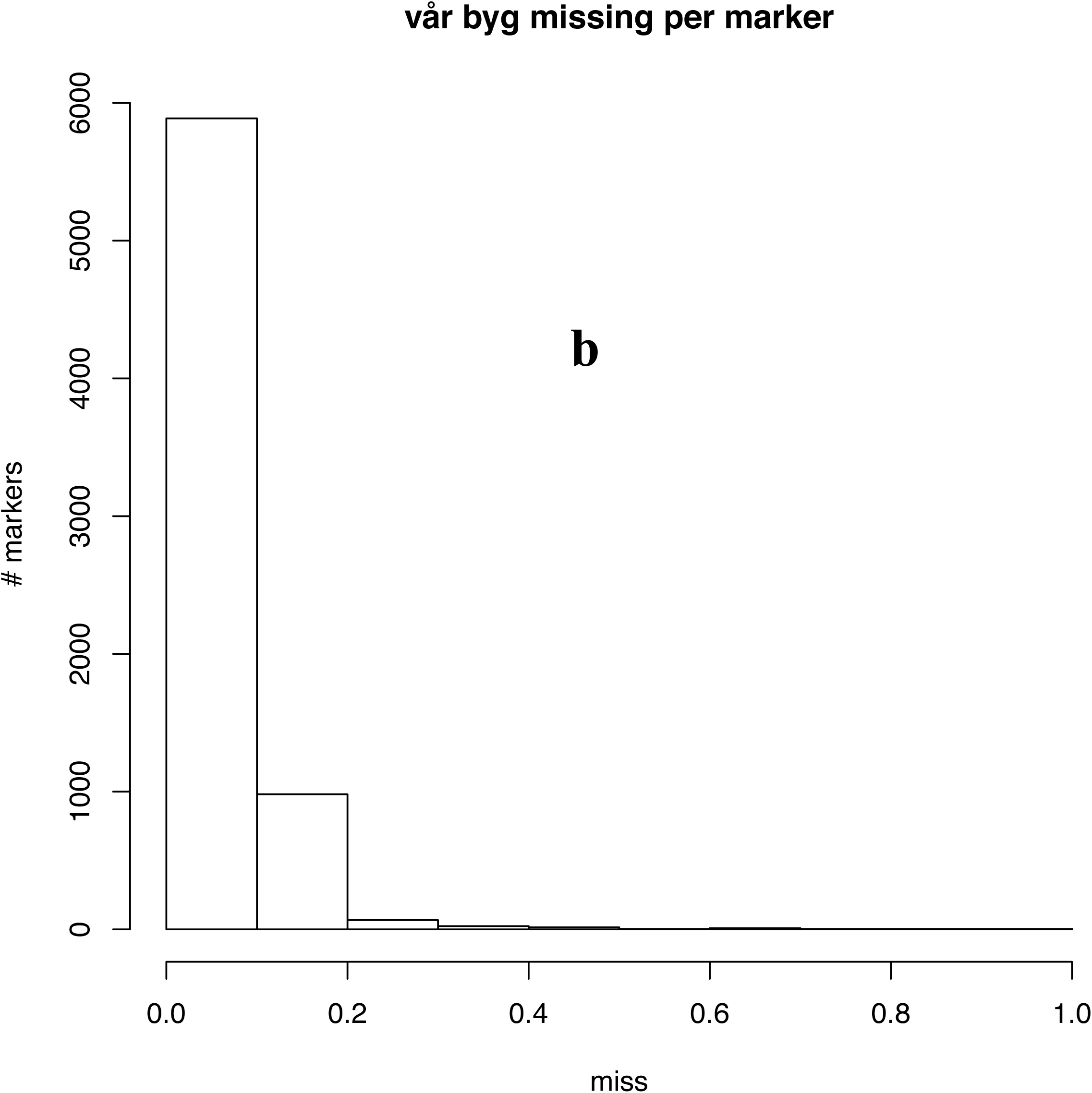
a: SNP markers MAF distribution before filtering. b: missing distribution of SNP markers before filtering.

### Population structure

Population structure among 1,329 Barley lines was investigated using principal component analysis on the genomic relationship matrix, G. The first two components were plotted, (Fig.3a), and each dot in the plot represent a line and the colors indicate the breeding cycles. The first two components explained 33 % and 12 % of the total markers variance, respectively. There was no clear grouping in the population even though the lines were derived from 4 different breeding cycles. To visualize the relationship among the lines, a heat map was produced (Fig.3b). Some lines were more related to each other than to others due to common ancestors but there were no strong sub-groups in the population.

**Fig.3.**
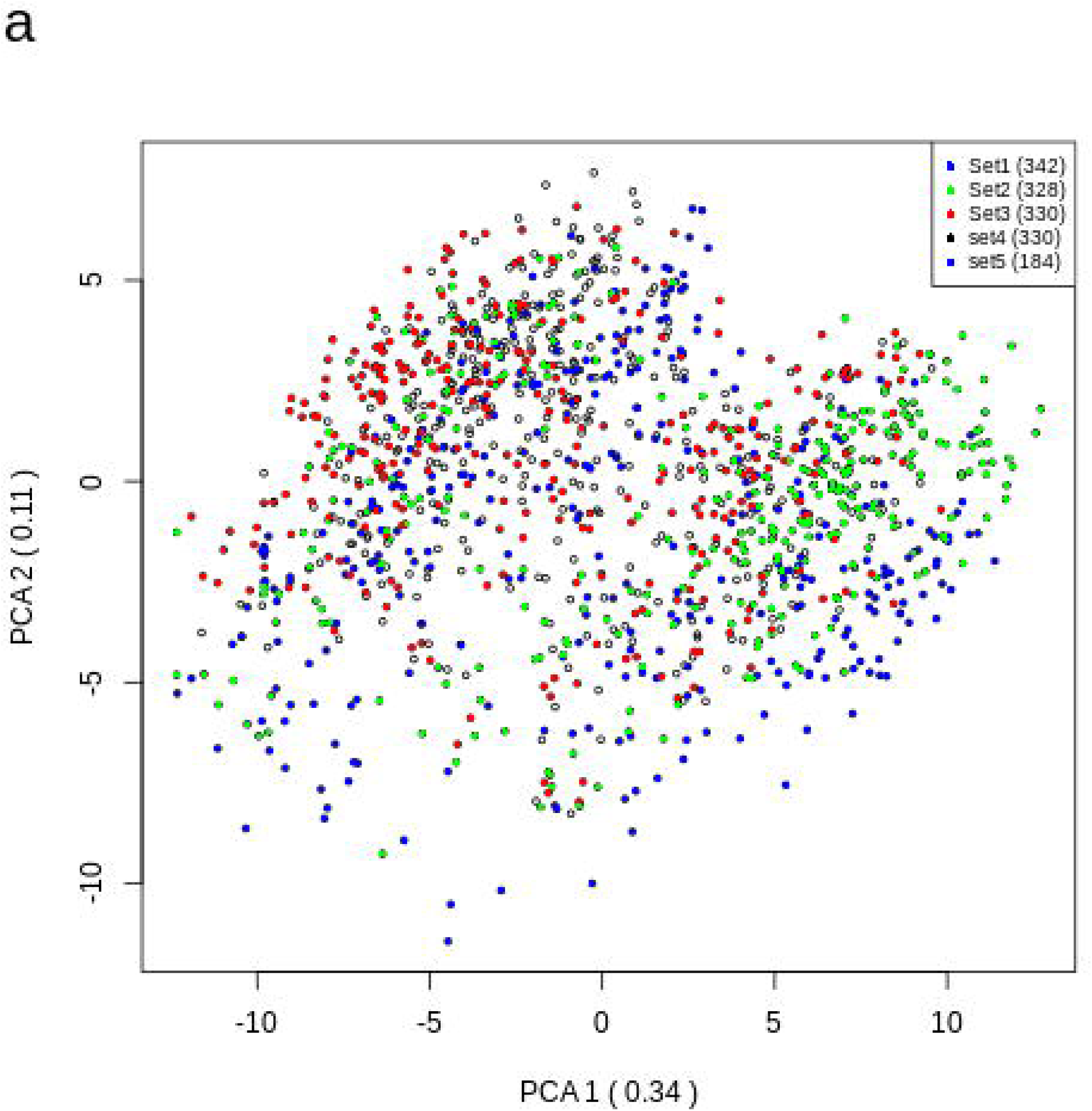
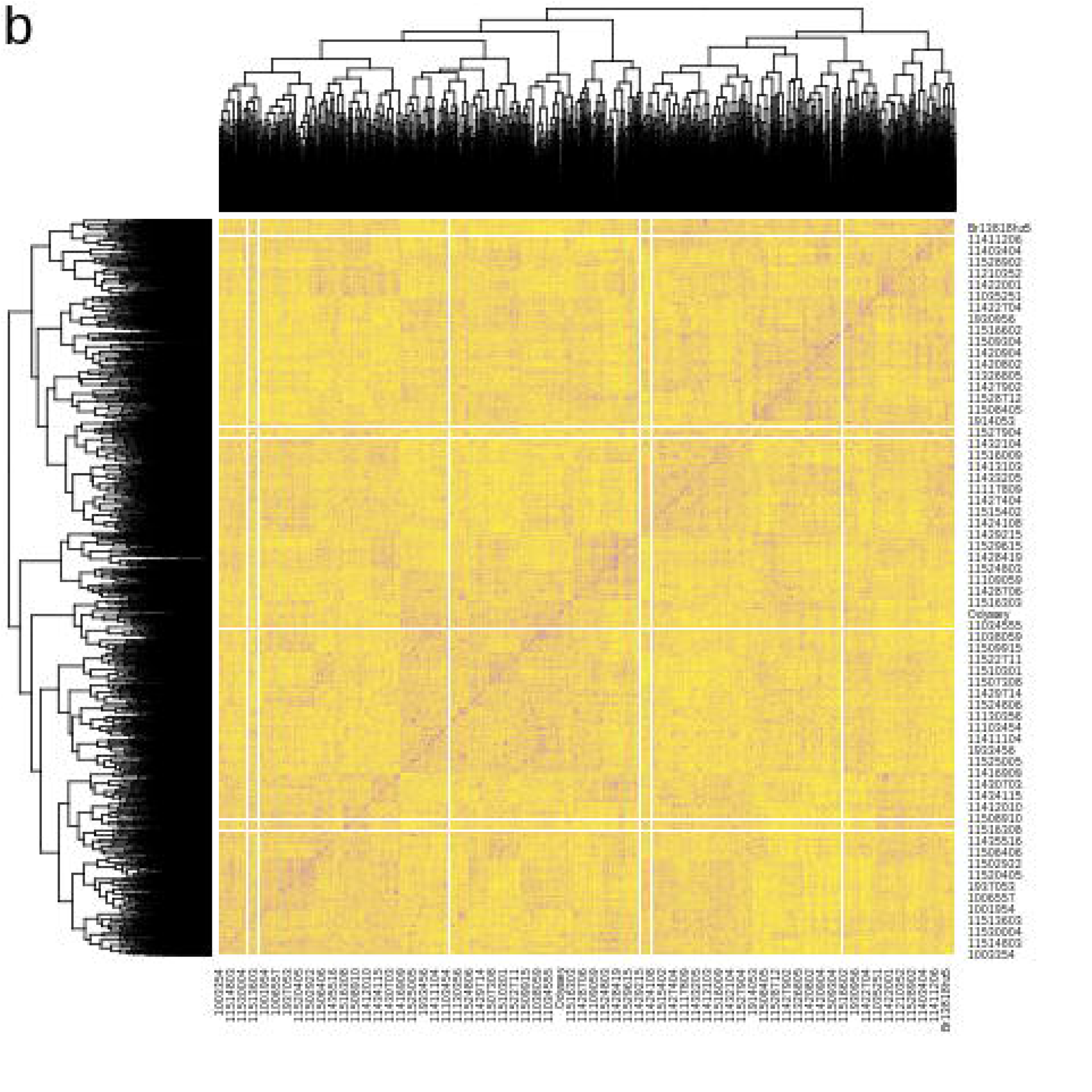
a: First two principal components of principal component analysis for the determination of population structure in barley lines. The colors represent different breeding cycles, b: Heatmap of relationship matrix (G).

### Heritability and predictive ability

Narrow sense heritability was estimated for each malting quality trait and can be found in Table 2. Heritability ranged from 0.31 for filtering speed to 0.65 for wort color. Most of the traits had a medium heritability.

**Table 2.**
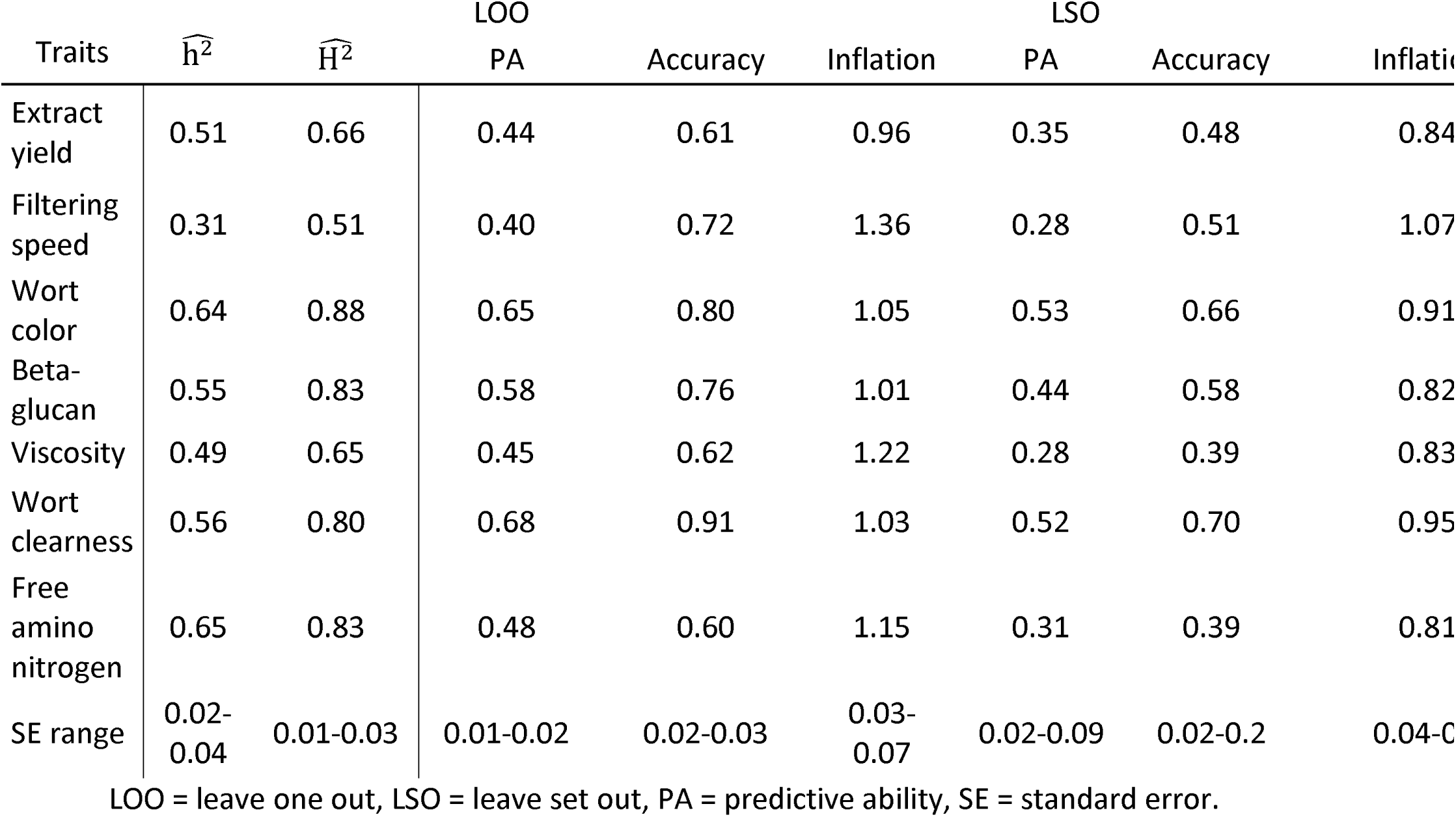
Estimates of heritability of malt quality traits, predictive ability, prediction accuracy, inflation, and their standard error for leave one out and leave set out cross-validation schemes.

Predictive ability for different malting quality traits for LOO and LSO cross-validation schemes can be found in Table 2. Predictive abilities were calculated as the correlation between the 1,329 lines GEBVs and the means of phenotypes corrected for effects of location, year and trial. Overall, as expected, LOO cross-validation method had a higher predictive ability than LSO and k-fold cross-validations methods. The highest predictive ability in LOO were for wort clearness and the lowest was for filtering speed with values of 0.40 and 0.68, respectively. In LSO method, the highest predictive ability was for wort color (0.53) and the lowest was for both filtering speed and viscosity (0.28). Moving from LOO method to LSO method, predictive ability from viscosity had the biggest loss (37.8 %) and wort color had the least lost with only 17.5 % on average the loss was 26.7% for the seven malt quality traits.

Prediction accuracy from three different fold cross-validation strategies for malting traits is shown in Fig.4. Among all the traits, wort clearness had the highest prediction accuracy. It was very close to the maximum value that, theoretically, can be achieved. Among the random cross-validation tests as expected, the tenfold cross-validation had the highest predictive accuracy. In all traits, the predictive accuracy increased by moving from twofold to fourfold cross-validation. For wort clearness, wort color, and beta-glucan, the increase in predictive accuracy increased a bit from fourfold to tenfold cross-validation as well. Comparing average accuracies from fourfold cross-validation sets (Fig.4, beta-glucan =0.72, extract yield= 0.59, free amino nitrogen = 0.56, viscosity = 0.63, wort clearness = 0.87, wort color = 0.76) to LSO accuracies (Table 2) it is clear that LSO accuracies are lower than the corresponding accuracies from fourfold cross-validation sets that had similar size of the training populations.

**Fig.4.**
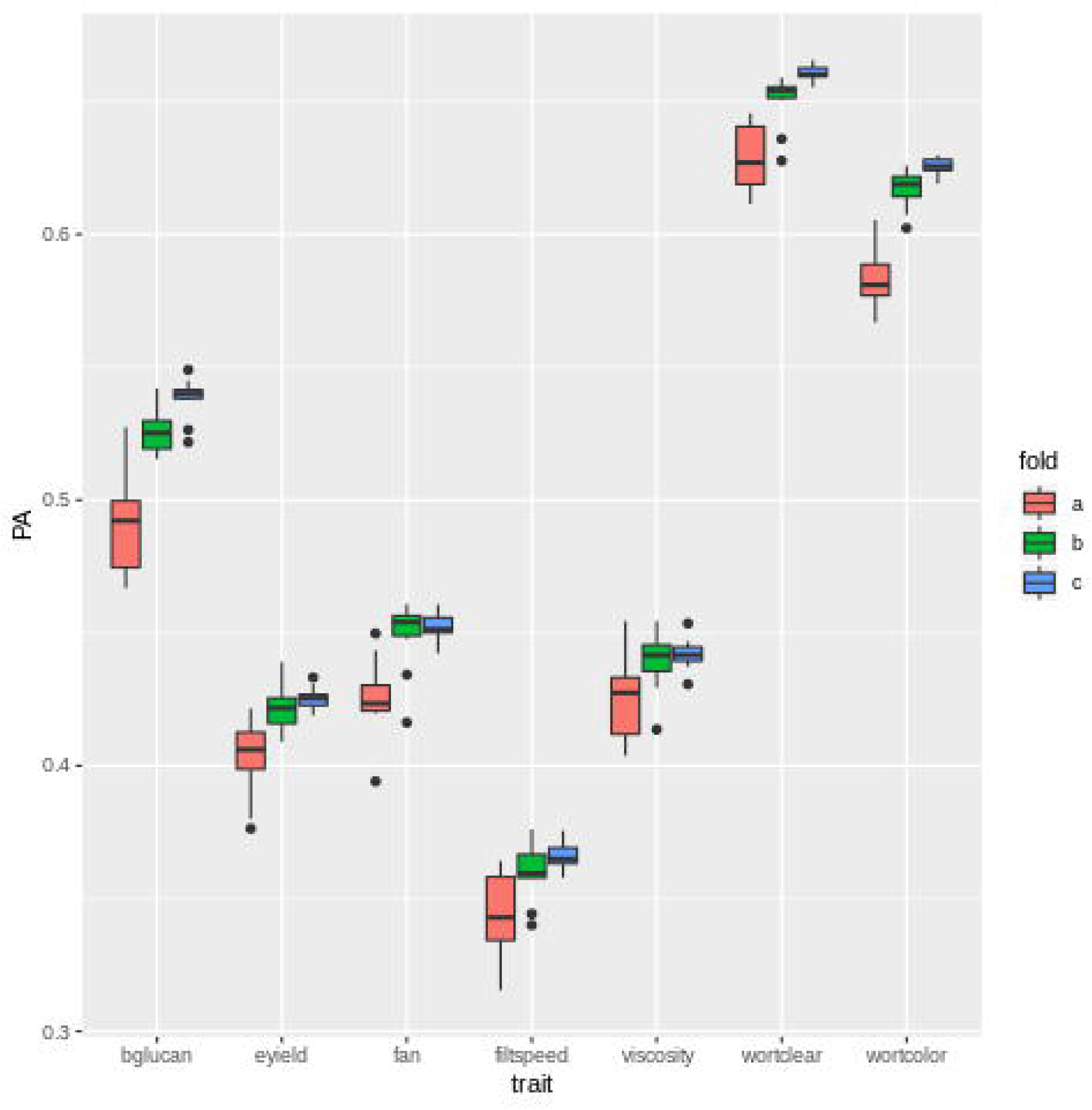
Predictive accuracy of various malting quality traits for two- (red), four- (green), and ten-fold cross-validation scheme (blue). The thick horizontal line is the median, the lower and upper hinges correspond to the 25th and 75th percentiles (first and third quartiles). The whiskers extends from the hinges 1.5 times the distance between the first and third quartiles, data points beyond the whiskers are plotted individually. Free amino nitrogen abbreviated to FAN.

Genetic correlation among the seven traits as well as the correlation among line effects, primarily capturing genetic variation that is not captured by additive effects of markers were calculated (Table 3). Ten of the trait combinations have unfavorable additive genetic correlations, and unfavorable correlations for the combined line and genomic effects. This means that it can be very difficult to select a line that is superior for all malt quality traits for either breeding or use as a commercial variety.

**Table 3.** Correlations among genetic effects of the seven malting quality traits. Above the diagonal the correlation among line + additive genetic effects, capturing broad sense genetic variation, are listed. Below the diagonal additive genetic correlations are listed.^1^

|  | Extract<br>yield | Filtering<br>speed | Wort color | Beta-glucan | Viscosity | Wort<br>clearness | Free amino<br>nitrogen |
| --- | --- | --- | --- | --- | --- | --- | --- |
| Extract<br>yield | - | -0.33 | 0.41 | -0.44 | -0.37 | -0.08 | 0.49 |
| Filtering<br>speed | -0.26 | - | -0.42 | 0.16 | -0.01 | -0.28 | -0.33 |
| Wort<br>color | 0.47 | -0.41 | - | -0.58 | -0.57 | 0.07 | 0.82 |
| Beta<br>glucan | -0.37 | 0.00 | -0.60 | - | 0.95 | 0.11 | -0.70 |
| Viscosity | -0.26 | -0.19 | -0.56 | 0.94 | - | 0.22 | -0.67 |
| Wort<br>clearness | -0.18 | -0.16 | -0.13 | 0.35 | 0.36 | - | -0.12 |
| Free<br>amino<br>nitrogen | 0.53 | -0.26 | 0.83 | -0.71 | -0.67 | -0.35 | - |

<sup>1</sup>Range of standard errors: 0.01-0.02

To illustrate the consequence of the genetic correlations among the malting quality traits, the best 200 advanced lines per trait based on genomic breeding values were selected and the overlapping lines were shown as a Venn diagram in Fig.5. The traits were divided into two groups based on the genetic correlations among traits. Fifteen lines were common among the top 200 lines for the first group of traits (extract yield, Beta glucan, viscosity, and wort clearness). In the second group of traits (wort color, free amino nitrogen, and filtering speed), only one line was common among all top 200 lines. The percentage of the lines that fall into top list for only one of the traits were between 6.5 to 23.9 in the first group of traits and 29.1 to 30.8 in the second group of traits.

**Fig.5.**
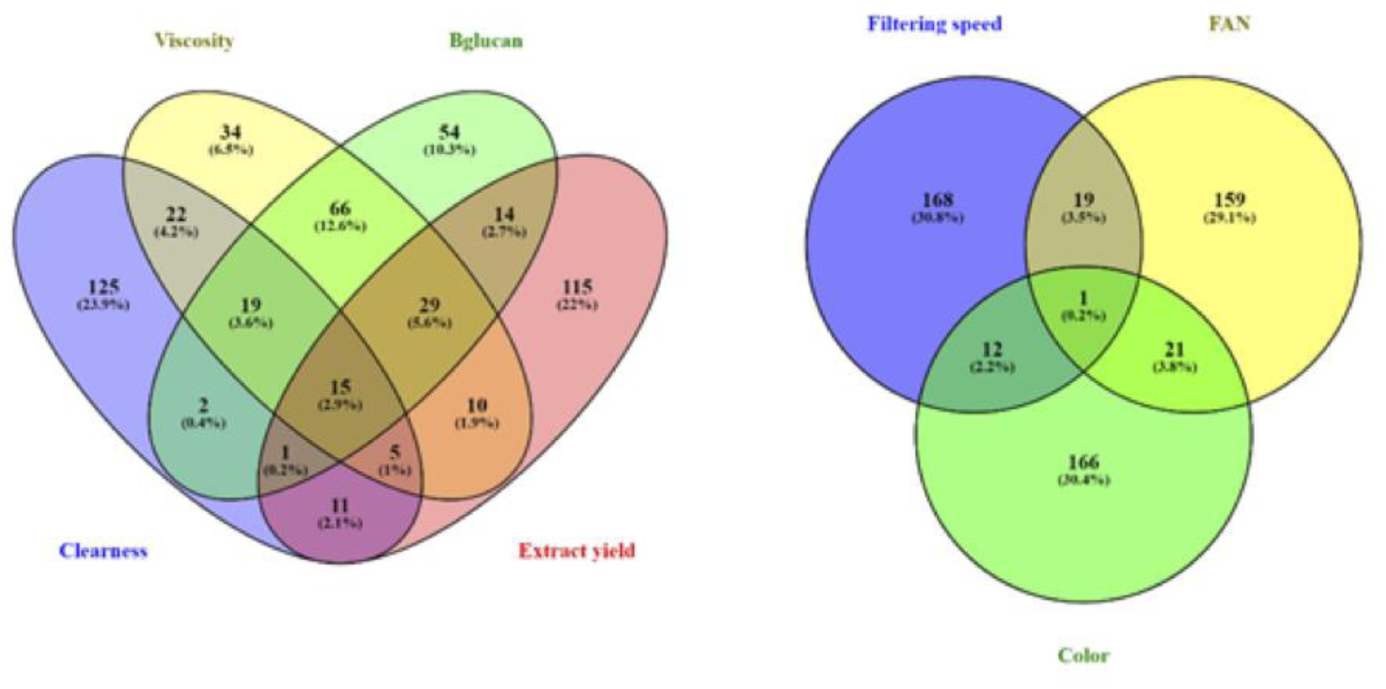
The dispersion of the best 200 lines for every malting quality based on genomic breeding values.

## Discussion

Malting quality traits are important in spring barley breeding as they are directly related to the amount and quality of brewed beer and other end products of malted barley. The identification and selection of the best malting barley lines is time consuming and expensive, since obtaining malting quality data is labor intensive and data on a large number of lines is needed to ensure sufficient selection intensity. Genomic selection can reduce the breeding cost since a lower number of lines need to be phenotyped to reach the same selection intensities, by using genomic prediction for the lines that are not being phenotypically assessed for malting quality. In addition, it can shorten the generation interval in breeding schemes by selecting lines in earlier stages of the breeding program. In this study, we explored the potential use of genomic selection in malting barley breeding. The population structure was analyzed to check for any strong sub-population and the ability to predict breeding values for malting quality were assessed for different scenarios using cross-validation.

The population consisted of material from four consecutive breeding cycles. Each year, a number of parents were crossed, and the best new lines were selected to be either new parents, become commercial varieties, or both. A few parents were present in more than one breeding cycle, but none of the parents was used in all four breeding cycles. The principle component analysis and heat map (Fig.3 a, b) showed strong genetic relationships among lines but no clear sup-populations in the material. This makes interpretation of results from different cross-validation schemes for genome wide selection less complicated than in a population with several subpopulations (Guo et al. 2014; Heslot et al. 2015; Sallam et al. 2015).

The estimates of narrow sense heritability 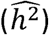 of different malting traits ranged between 0.31 for filtering speed and 0.65 for wort color, the latter is comparable to the 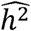 of wheat quality traits in Kristensen et al. (2019). For extract yield, the broad sense heritability 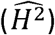 was 0.66 which was lower than the 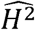 found in a previous study by Schmidt et.al. (2015) 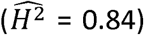 but comparable to the broad sense heritability reported by Bhatta et al. (2020) 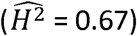. For the other three traits that are in common between these studies, our broad sense heritability estimates are comparable to the ones reported by Schmidt *et al.* (2015) and higher than reported by Bhatta *et al.* (2020). For free amino acid, beta-glucan, and viscosity 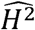 estimates were 0.79, 0.85, and 0.68, respectively (Schmidt et al. 2015) and free amino acids and beta-glucan 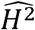 were 0.74 and 0.66, respectively (Bhatta et al. 2020). As expected, all reported 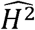are larger than the 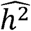. H^2^ is the proportion of phenotypic variation that is due to total genetic variation including both additive and non-additive genetic effects (such as epistatic effects). As h^2^ is the proportion of phenotypic variation that is only additive genetic variation, it will often be considerably smaller than H^2^. The genomic prediction models used in the current study is based on the additive genetic effects and not the epistatic effects, h^2^ can tell us how much of the phenotypic variation we at best can expect to be able to predict using standard genomic prediction methods. The additive effect is also the effect that is most important in selecting new parents in the breeding program. Therefore, h^2^ are most relevant in this context.

There are a number of factors affecting the predictive ability of a genomic prediction model. The heritability is one of these factors (Sallam et al. 2015; Duangjit et al. 2016; Lozada and Carter 2019). As expected, we found that traits with higher h^2^ had higher accuracy of genomic prediction. Another factor is the size of the training population (Berro et al. 2019), we investigated this by three different CV strategies, where 50%, 75%, or 90% of the lines were used as the training population corresponding to 665, 997, and 1196 lines in the training population, respectively. Most traits showed little increase in predictive ability between the fourfold and tenfold cross-validation schemes indicating that the 997 lines in the fourfold CV strategy had enough information to ensure a robust estimation of population parameters in the genomic model. Thus, the number of lines in 3 breeding cycles could be sufficient for an operational training population, and accuracies would not be expected to increase further for most traits by increasing the number of lines in the training population. However, accuracies of genomic breeding values of wort clearness, wort color and Beta-glucan might increase further with a larger training population (Fig.4). It is worth noting that the genomic relationships were less strong across sets than they were across the random fourfold validation sets. Therefore, prediction across sets will have lower accuracies than estimated by the fourfold cross-validation sets.

A third factor that affects the prediction accuracy of genomic prediction is the strength of the genomic relationship between the lines in the validation population and the lines in the training population. In our study, the 1,329 lines resulted in a robust and trustworthy genomic prediction within breeding cycle. The accuracy using the LOO cross-validation strategy ranged from 0.60 for free amino nitrogen to 0.91 for wort clearness. Overall, in LOO strategy, all different types of relatedness present in the population was considered and exploited in the prediction model. When a line was used for validation, there would often be full or half sibs of that line in the training population. In addition, the LOO strategy mimics the situation in a breeding scheme, where some of the lines in each breeding cycle have own phenotypes and others do not *i.e*. a situation with a yearly update of the training population.

In a commercial breeding program, every year new crosses are made between a set of elite parent lines, and the resulting progenies are advanced to select the best breeding lines for registration. Some selection could be conducted before any of the lines in the breeding cycle have their own malting quality phenotype. This situation is mimicked in the LSO cross-validation strategy. As expected, the LSO predictive ability in general was less than LOO strategy. Nielsen et al. (2016) found similar results in their study on seed quality traits and they concluded the reason was a smaller training population size in LSO, where a quarter of the lines in the study is in the validation population compared to LOO strategies. However, as most traits in this study showed little increase in predictive ability between the four-fold and ten-fold cross-validation, lower genomic relationship between training and validation population in LSO validation probably contributed considerably to the difference in predictive ability between LOO and LSO. This interpretation of the results was supported by the lower accuracy in LSO tests compared to fourfold cross-validation sets. The decrease in predictive abilities in LSO compared to the LOO strategy varies for different traits. The biggest decrease was in filtering speed where the accuracy was reduced by 0.23. The trait with smallest difference between LSO and LOO was extract yield, which was reduced by 0.13.

Looking at the accuracies of the LOO validation set in Table 2, we can divide the seven traits into three groups according to how well the genomic model perform. For the first group, comprising of wort clearness only, the model performs very well with no significant scope for improvement. This performance is similar to what we see in genomic prediction of baking quality traits in wheat (Kristensen et al. 2019). For the second group, comprised by wort color, filtering speed, and beta-glucan, the accuracies in the LOO CV scheme were in the range of 0.85-0.88. As discussed above, for the two latter traits accuracies could be improved, when new lines are added to the training population. For the third group, comprised by extract yield, viscosity, and free amino nitrogen, the accuracies in the LOO CV scheme were in the range of 0.60-0.62. Fig.4 shows that increasing training population size only resulted in small improvements in PA for this group. The same tendency is seen in prediction of wheat quality traits, when the training population exceeds 400 lines (Kristensen et al. 2019). An alternative route to increased accuracy is to analyze genetically correlated traits in a multi-trait genomic prediction model (Bhatta et al. 2020). Extract yield and viscosity are two of the three most important malting quality traits (the third being beta-glucan), and thus it would be relevant to investigate models and methods that can improve accuracy of genomic prediction in this third group of malting quality traits in future studies.

The narrow sense heritabilities in Table 2 combined with the phenotypic variance seen in Fig 1 indicated that there is ample genetic variation available for selection. For some traits, high breeding value is best but for other traits the lowest breeding value is desirable and for the remaining traits the medium range values are optimal. For example, lines that produce more extract yield are desirable for brewing companies so the higher value in genomic predictive value is better. On the other hand, having lots of beta-glucan in the malt will slow down the enzymes activity and increase the viscosity that is not considered as a desirable character and the lines with lower estimated breeding value should be selected. For free amino nitrogen, a value close to the population mean average is best and lines with breeding values around zero should be selected. As can be seen from Table 3, lines with desirable high breeding values for extract yield will tend to have undesirable high values of free amino nitrogen. Thus, if breeders select heavily for extract yield, they would unfavorably select for increased free amino nitrogen. Proper multivariate selection criteria need to be developed. A further increase in prediction accuracy can possibly be obtained by the use of multi-trait genomic selection with each trait weighted by their economic importance in the breeding program (Akdemir et al. 2019).

## Conclusions

Predictive ability of malting quality traits in spring barley shows that genomic prediction both within and across breeding cycles can be applied in practical breeding programs, where the main target is selecting high merit malting barley varieties. The use of a homogenous commercial population did not hamper the efficient use of genomic selection. The predictive ability when moving from one breeding cycle to the next is acceptable and can be exploited in commercial breeding programs. Selecting the best line with all desirable characteristic for malting quality traits is still difficult using single trait genomic prediction though.

## Declarations

### Author Contributions

Conceptualization, PS, JJ, JDJ, VE, and AJ.; Data curation, NHK, VE and JO; Formal analysis, VE and PS; Funding acquisition, NHK, JJ, JDJ and AJ; Investigation, VE, NHK and PS; Methodology, VE, NHK, PS, JJ, and AJ; Project administration, JJ and AJ; Resources, PS, VE, NHK, JDJ, and JO; Software, VE, PS and JJ; Supervision, JJ, JDJ and AJ; Validation, PS and VE; Visualization, VE, and PS; Writing—original draft, VE; Writing—review & editing, PS, JJ, JDJ, NHK, JO and AJ

## Acknowledgments

We would like to thank Hanne Svenstrup and Jette Andersen from Nordic Seed A/S, Denmark for supporting laboratory work and phenotyping.

## Availability of data and material

Data will be made available as supplementary material upon acceptance of the MS.

## Code availability

Not applicable

## References

Akdemir D, Beavis W, Fritsche-Neto R, et al (2019) Multi-objective optimized genomic breeding strategies for sustainable food improvement. Heredity (Edinb) 122:672–683. https://doi.org/10.1038/s41437-018-0147-1

Bamforth CW (2009) Current perspectives on the role of enzymes in brewing. J Cereal Sci 50:353–357

Bamforth CW (2003) Barley and malt starch in brewing: a general review. Tech Q Master Brew Assoc Am 40:89–97

Berro I, Lado B, Nalin RS, et al (2019) Training Population Optimization for Genomic Selection. Plant Genome 12:190028. https://doi.org/10.3835/plantgenome2019.04.0028

Bhatta M, Gutierrez L, Cammarota L, et al (2020) Multi-trait Genomic Prediction Model Increased the Predictive Ability for Agronomic and Malting Quality Traits in Barley (Hordeum vulgare L.). G3 Genes|Genomes|Genetics g3.400968.2019. https://doi.org/10.1534/g3.119.400968

Celus I, Brijs K, Delcour JA (2006) The effects of malting and mashing on barley protein extractability. J Cereal Sci 44:203–211

Cericola F, Jahoor A, Orabi J, et al (2017) Optimizing Training Population Size and Genotyping Strategy for Genomic Prediction Using Association Study Results and Pedigree Information. A Case of Study in Advanced Wheat Breeding Lines. PLoS One 12:e0169606

Delcour JA, Verschaeve SG (1987) Malt diastatic activity. Part II. A modified EBC diastatic power assay for the selective estimation of beta-amylase activity, time and temperature dependence of the release of reducing sugars. J Inst Brew 93:296–301

Duangjit J, Causse M, Sauvage C (2016) Efficiency of genomic selection for tomato fruit quality. Mol Breed 36:29. https://doi.org/10.1007/s11032-016-0453-3

Evans DE, Collins H, Eglinton J, Wilhelmson A (2005) Assessing the Impact of the Level of Diastatic Power Enzymes and Their Thermostability on the Hydrolysis of Starch during Wort Production to Predict Malt Fermentability1. J Am Soc Brew Chem 63:185–198

Fernando RL, Habier D, Stricker C, et al (2008) Genomic selection. Acta Agric Scand Sect A 57:192–195

Gao W, Clancy JA, Han F, et al (2004) Fine mapping of a malting-quality QTL complex near the chromosome 4H S telomere in barley. Theor Appl Genet 109:750–760

Goddard ME, Hayes BJ (2007) Genomic selection. J Anim Breed Genet 124:323–330

Guo G, Zhao F, Wang Y, et al (2014) Comparison of single-trait and multiple-trait genomic prediction models. BMC Genet 15:1–7

Hayes BJ, Bowman PJ, Chamberlain AC, et al (2009) Accuracy of genomic breeding values in multi-breed dairy cattle populations. Genet Sel Evol 41:51

Hayes PM, Liu BH, Knapp SJ, et al (1993) Quantitative trait locus effects and environmental interaction in a sample of North American barley germ plasm. Theor Appl Genet 87:392–401

Heffner EL, Sorrells ME, Jannink J-L (2009) Genomic Selection for Crop Improvement. Crop Sci 49:1–12

Heslot N, Jannink J-L, Sorrells ME (2015) Perspectives for genomic selection applications and research in plants. Crop Sci 55:1–12. https://doi.org/10.2135/cropsci2014.03.0249

Jannink JL, Lorenz AJ (2010) Genomic selection in plant breeding: from theory to practice

Kristensen PS, Jensen J, Andersen JR, et al (2019) Genomic prediction and genome-wide association studies of flour yield and alveograph quality traits using advanced winter wheat breeding material. Genes (Basel) 10:1–19. https://doi.org/doi:10.3390/genes10090669

Li CD, Cakir M, Lance R (2009) Genetic Improvement of Malting Quality through Conventional Breeding and Marker-assisted Selection. In: Genetics and Improvement of Barley Malt Quality. Springer, Berlin, Heidelberg, Berlin, Heidelberg, pp 260–292

Lorenz AJ, Chao S, Asoro FG, et al (2011) Genomic Selection in Plant Breeding: Knowledge and Prospects. Adv Agron 110:77–123

Lozada DN, Carter AH (2019) Accuracy of Single and Multi-Trait Genomic Prediction Models for Grain Yield in US Pacific Northwest Winter Wheat. Crop Breeding, Genet Genomics 1:e190012. https://doi.org/10.20900/cbgg20190012

Madsen P, Jensen J (2000) DMU

Meuwissen TH, Hayes BJ, Goddard ME (2001) Prediction of total genetic value using genome-wide dense marker maps. Genetics 157:1819–1829

Nielsen NH, Jahoor A, Jensen JD, et al (2016) Genomic Prediction of Seed Quality Traits Using Advanced Barley Breeding Lines. PLoS One 11:e0164494

Qi J, Chen J, Wang J, et al (2005) Protein and hordein fraction content in barley seeds as affected by sowing date and their relations to malting quality. J Zhejiang Univ Sci B 6:1069–1075

Riedelsheimer C, Melchinger AE (2013) Optimizing the allocation of resources for genomic selection in one breeding cycle. Theor Appl Genet 126:2835–2848

Saghai-Maroof MA, Soliman KM, Jorgensen RA, Allard RW (1984) Ribosomal DNA spacer-length polymorphisms in barley: mendelian inheritance, chromosomal location, and population dynamics. Proc Natl Acad Sci U S A 81:8014–8018

Sallam AH, Endelman JB, Jannink J-L, Smith KP (2015) Assessing Genomic Selection Prediction Accuracy in a Dynamic Barley Breeding Population. Plant Genome 8:plantgenome2014.05.0020. https://doi.org/10.3835/plantgenome2014.05.0020

Schmidt M, Kollers S, Maasberg-Prelle A, et al (2015) Prediction of malting quality traits in barley based on genome-wide marker data to assess the potential of genomic selection. Theor Appl Genet 129:203–213

VanRaden PM (2008) Efficient Methods to Compute Genomic Predictions. J Dairy Sci 91:4414–4423

